# Does a history of sexual and physical childhood abuse contribute to HIV infection risk in adulthood? A study among post-natal women in Harare, Zimbabwe

**DOI:** 10.1101/333872

**Authors:** Simukai Shamu, Patience Shamu, Christina Zarowsky, Marleen Temmerman, Tamara Shefer, Naeemah Abrahams

## Abstract

**Background:** Sexual and physical abuse in childhood creates a great health burden including on mental and reproductive health. A possible link between child abuse and HIV infection has increasingly attracted attention. This paper investigated whether a history of child physical and sexual abuse is associated with HIV infection among adult women.

**Methods:** A cross sectional survey was conducted among 2042 postnatal women (mean age=26y) attending six public primary health care clinics in Harare, Zimbabwe within 6 weeks post-delivery. Clinic records were reviewed for mother’s antenatal HIV status. Participants were interviewed about childhood abuse including physical or sexual abuse before 15 years of age, forced first sex before 16, HIV risk factors such as age difference at first sex before age 16. Multivariate analyses assessed the associations between mother’s HIV status and child physical and sexual abuse while controlling for confounding variables.

**Results:** More than one in four (26.6%) reported abuse before the age of 15: 14.6% physical abuse and 9.1% sexual abuse,14.3% reported forced first sex and 9.0% first sex before 16 with someone 5+ years older. Fifteen percent of women tested HIV positive during the recent antenatal care visit. In multivariate analysis, childhood physical abuse (aOR 3.30 95%CI 1.58- 6.90), sexual abuse (3.18 95%CI: 1.64-6.19), forced first sex (aOR 1.42, 95%CI: 1.00-2.02), and 5+ years age difference with first sex partner (aOR 1.66 95%CI 1.09-2.53) were independently associated with HIV infection.

**Conclusion:** This study confirms that child physical and/or sexual abuse increases risk for HIV acquisition. Further research is needed to assess the pathways to HIV acquisition from childhood to adulthood. Prevention of child abuse must form part of the HIV prevention agenda in Sub-Saharan Africa.

## Introduction

Child abuse is a violation of the fundamental rights of children. Although research on child abuse is growing globally, research on childhood exposure to violence is a neglected area in many parts of Africa. Globally, 180/1000 girls are abused sexually and 163/1000 are abused physically^1^. An extensive review of studies in Africa reported among the highest prevalence of child abuse globally^2^. The review revealed that both boys and girls are affected by abuse. The prevalence of physical or sexual violence against girls in Africa ranges between 2% and 64%^2^ and the wide range is a result of the various definitions of violence adopted and settings where the study was conducted (see also http://www.knowviolenceinchildhood.org/). Most of the abuse of children happens at home or in school^3^. Drivers of child abuse include family disorganisation and dysfunction^2,4^ often related to contexts of poverty and structural violence, cultural factors and community norms accepting violence such as bullying^5^, legal frameworks that allow the use of violence to discipline children^6^ and gender inequity and inequality^7,8^. For example, in Zimbabwe the use of corporal punishment in schools is permitted to discipline children under special circumstances - heads of schools exercise corporal punishment while teachers can obtain permission to use corporal punishment from the school head. In each instance where corporal punishment is used a register is kept (Secretary of Education Circular P 35, 1993; Statutory Instrument 65 of the Constitution of Zimbabwe, 1992). Nevertheless, corporal punishment is used outside of these conditions as it goes undocumented, goes beyond disciplining to set limits and having detrimental effects to children ^6^. Research has also highlighted the absence of alternative ways of disciplining children leading to the use of violence both in schools and at home^3,8^. Shumba argues that the use of violence against children in Zimbabwean schools is an extension of violent ways of disciplining a child that teachers transfer from their roles as parents and adults in the community to the school^6^.

The few studies on sexual violence in Zimbabwe show that sexual violence against girls is high^9^ while more than one in five (21.6%) report forced first sexual intercourse. The malfunctioning of the family institution to protect or look after children, orphanhood, maternal challenges and non-attendance of school were found to be associated with sexual abuse in Zimbabwe^9^. Studies in high income countries have shown that abuse in childhood is likely to result in later life challenges including mental health problems (substance use, suicide behaviour, depression, stress) and sexually transmitted infections ^10–13^. More literature is gathering evidence on the relationship with HIV^5^ and these include a few studies being conducted in Sub-Saharan Africa^8,9^. Despite this, the evidence base is still thin and no studies have assessed multiple childhood adversities and their association with HIV in pregnancy in one study. Dedicated research on sexual violence is seldom a focus in Zimbabwe as child sexual abuse is researched as part of other studies^16^. The intersection of age with gender and other inequalities facilitate HIV vulnerability. Women’s sexual relationships with older men in Zimbabwe have been reported to increase HIV incidence^17^. Having sexual intercourse with a minor who is five years younger than the person having sex with them is also considered a legally abusive act as it indicates possible coercive or abusive experiences. Research shows an association between age and unwanted sexual practices ^17^ which increase the risk of HIV infection.

This study provides an opportunity to assess if child physical/sexual abuse is associated with HIV diagnosed in pregnancy. We used data in a women’s health research project examining intimate partner violence and HIV diagnosed during pregnancy to estimate here the prevalence of childhood adversities (child sexual abuse, child physical abuse, forced first sex and first sex with older men) and to assess the relationship between these childhood adversities with HIV infection. We tested the hypothesis that a woman’s history of childhood adversities contributes to her HIV positive status in antenatal care.

## Methods

We conducted a cross sectional study in Harare, Zimbabwe in 2011 among 2042 women aged 18-49 years attending postnatal care at six public primary health care facilities in resource limited formal urban locations. We reported detailed methodology for the study elsewhere^18^. Face to face interviewer administered structured interviews were conducted by trained female fieldworkers to collect demographic and childhood abuse experiences data for this study. HIV status information were abstracted from clinic records. Women were recruited while queuing for postnatal services and baby immunisation sessions at the six clinics. Interviews were conducted in privacy. The study clinics had both antenatal and postnatal services and were owned and managed by the Harare City Council.

We measured child physical abuse by asking each participant whether anyone ever “excessively” beat or physically mistreated the woman in any way before age 15. We prefixed the word “excessive” to categorise abuse differently from culturally acceptable ways of enforcing discipline in Zimbabwe which were common in the study setting^6^. Any response in the affirmative was considered as childhood physical abuse. For child sexual abuse we followed the WHO Multi-country study on women’s health and domestic violence^19^ by asking whether anyone ever forced a woman to have sex or to perform a sexual act or ever touched her sexually when she did not want to before she turned age 15. Any response in the affirmative was recorded as childhood sexual abuse. To assess forced first sexual intercourse respondents were asked whether their first sexual intercourse occurred when they were willing, tricked, persuaded, forced or raped^19,20^. If a participant responded that they were forced or raped we defined it as forced first sexual intercourse. Participants who responded affirmatively and also reported first sex before age 16 were considered as having had forced first sex before age 16. We used this conceptualisation of forced first sex before age 16 in this analysis.

To assess first sex with someone 5+ years older while under 16 years we first asked women about their age and the age of their partners. The two ages were then compared to identify the age difference. Women were also asked the year in which they first had sexual intercourse. We constructed a variable that describes those who had sexual intercourse before age 16 and had first sex with someone five years or older than them. Although many studies have reported that age 18 was the median age of sexual debut of many women in Zimbabwe^21^ we used age 16 as the cut off to define childhood sexual abuse following the Sexual Offenses Act of Zimbabwe^22^. Intimate partner physical/sexual violence was measured using the WHO questionnaire on measuring domestic violence against women and girls^23^. The interviews also collected demographic information of the participants including education, employment status, marital status, and current or most recent partner’s demographic characteristics. HIV was tested through rapid test kits^18^. Health facilities in the study used the Determine^(tm)^ rapid test and positive results confirmed using Capillus, and the Western blot was used to resolve any conflicts. HIV results were abstracted from the patient records with permission from the City Health Department and the participant. This paper focuses on participants with available HIV test results.

We analysed data in Stata version 13 ^24^. Variable frequencies were calculated and presented in tables. Sociodemographic and childhood abuse experiences were described by HIV status using chi-square tests to illustrate statistical differences between groups. We reported statistical differences at 95% confidence levels (CI). Univariate analyses were performed before building a multiple regression model of factors associated with testing HIV positive. We performed a multivariate regression analysis with HIV as our outcome variable using a backward elimination process. We reported regression model output as adjusted Odds Ratios (aOR) with 95% CI and p values. The model controlled for known HIV covariates such as demographic variables (age, marital status and education), and history of intimate partner violence based on our knowledge of determinants of HIV in generalised epidemics such as in Zimbabwe.

The study was approved for ethics by the Medical Research Council of Zimbabwe and the University of the Western Cape Research Ethics Committees. The Harare City Health Department gave permission for the study to be conducted at the six clinics. All women were given contact details of organisations that provide counselling services for abused women for assistance as per the WHO guidelines for researching violence against women and girls^25^. We sought and received informed written consent for all participants while those mothers who were under 18 years of age provided informed written assent after parental/guardian consent was provided.

## Findings

We approached 2101 women to participate in the study and 2042 (97%) agreed and completed the interviews. A total of 1951 (95.5%) of those interviewed had available HIV test results in the clinic records. Demographic characteristics of participants (age, education, partners’ age and education, marital status, employment status, gravida and age differences with partners) with and without HIV test results did not differ (p>0.05) (data not shown).

Table 1 shows the participants’ demographic characteristics. Women were on average 26.4 years old [95% Confidence interval (CI) 26.17-26.68], over 9 in 10 (92.7%) attended at least secondary education. Close to a third (30%) were employed, 88.2% were married. Mean gravidity was 2.2. Participant’s partners were on average 31.3 years of age, that is, five years older than them. They were also more educated than them with 97% having completed at least secondary education (11 years of formal education).

**Table 1:** Socio-demographic characteristics of the sample (N=2042)

|  | **n** | **%** | **LCI** | **UCI** |
| --- | --- | --- | --- | --- |
| Age (mean) |  | 26.4 | 26.17 | 26.68 |
| Education: at least secondary | 1889/2037 | 92.7 |  |  |
| Woman employed | 606/2027 | 29.9 |  |  |
| Not married | 240/2041 | 11.8 |  |  |
| Gravida (mean) |  | 2.2 | 2.16 | 2.27 |
| Partner has at least secondary education | 1973/2030 | 97.2 |  |  |
| Partner's age (mean) |  | 31.3 | 31.04 | 31.61 |

Table 2 shows the prevalence of participants’ childhood sexual or physical abuse by HIV status. Fourteen percent (14.3%) of the women reported forced first sexual intercourse. Women in the sample generally partnered with men older than them. The mean age difference between women and their partners was 5.17 years and almost a tenth of the women (9.0%) reported having sex for the first time with someone five or more years older than them while they were below 16 years of age. Slightly more than a quarter (26.1%) of the women reported any form of childhood abuse before age 15. Almost one in ten (9.1%) women reported physical abuse before their 15^th^ birthday while 14.6% reported sexual abuse before age 15. Significant differences were found among all the four experiences (forced first sex, child physical abuse, child sexual abuse and age difference between the woman and her partner at first sex) and HIV status. More women who reported these experiences tested HIV positive than women who did not report these childhood abusive experiences and HIV risk factors.

**Table 2:** Prevalence of childhood abuse by HIV status (N=1945)

|  | Sample |  | HIV+ (n=299) |  | HIV-(n=1646) |  |  |
| --- | --- | --- | --- | --- | --- | --- | --- |
|  | % | CI | n | % | n | % | p-value |
| Forced first sexual intercourse | 14.3 | 12.81-15.85 | 58/299 | 19.4 | 218/1647 | 13.2 | 0.005 |
| 5+ year age difference at 1<sup>st</sup> intercourse before age 16 | 9.0 | 7.75-10.24 | 43/295 | 14.6 | 130/1649 | 7.8 | <0.0001 |
| Child sexual abuse | 14.6 | 13.05-16.19 | 46/299 | 15.4 | 131/1643 | 8.0 | <0.0001 |
| Child physical abuse | 9.1 | 7.83-10.39 | 57/299 | 19.3 | 227/1646 | 13.8 | 0.014 |
| **Adult experiences** |  |  |  |  |  |  |  |
| HIV positivity | 15.3 | 13.72-16.92 |  |  |  |  |  |
| Last 12mo physical/sexual IPV | 47.0 | 43.76-48.18 |  |  |  |  |  |

Fifteen percent (15.3%) of the women tested HIV positive. About 47% of the women reported experiencing physical or sexual intimate partner violence (IPV) in the 12 months preceding the most recent pregnancy.

Table 3 shows the results of multiple regression analysis that assessed factors associated with testing HIV positive in antenatal care. Childhood physical abuse (aOR 3.30 95%CI 1.58- 6.90), sexual abuse (3.18 95%CI: 1.64-6.19), forced first sex (aOR 1.42, 95%CI: 1.00-2.02), and 5+ years age difference with first sex partner (aOR 1.66 95%CI 1.09-2.53) were associated with HIV infection. The strongest associations (at least aOR 3.18) were between HIV and child sexual abuse and physical abuse. Women who were older than 24, not married, being less educated and having a partner at least 30 years old had higher odds of being HIV infected.

**Table 3:**
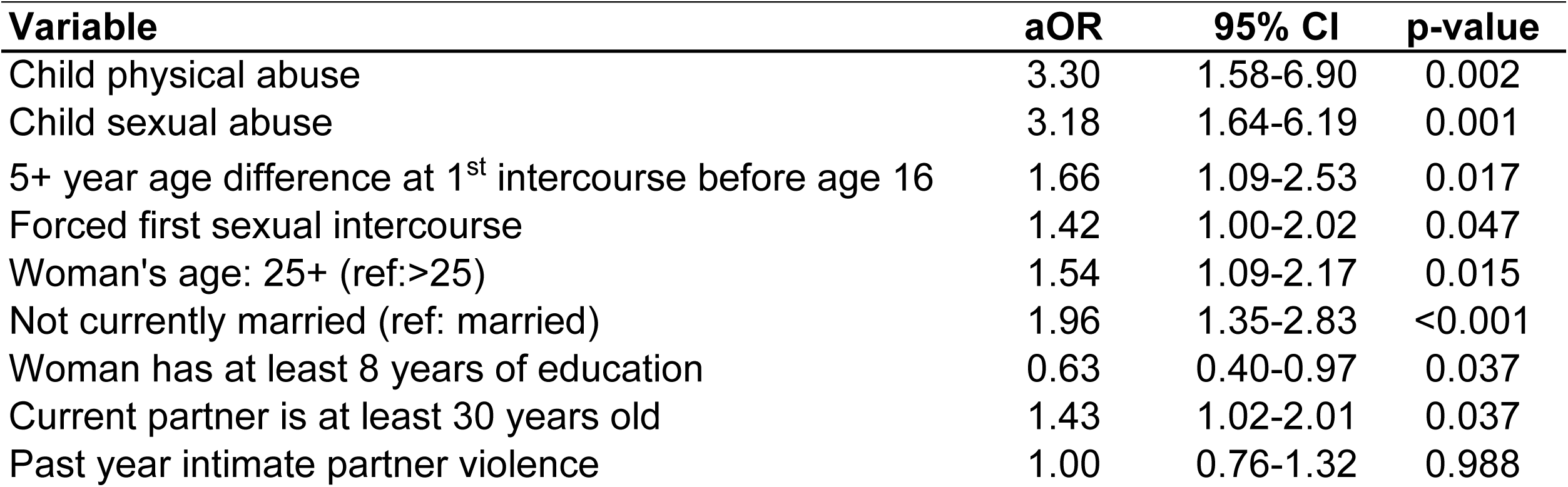
Multivariate logistic regression showing factors associated with HIV after adjusting for demographic factors and partner violence.

| **Variable** | **aOR** | **95% CI** | **p-value** |
| --- | --- | --- | --- |
| Child physical abuse | 3.30 | 1.58-6.90 | 0.002 |
| Child sexual abuse | 3.18 | 1.64-6.19 | 0.001 |
| 5+ year age difference at 1<sup>st</sup> intercourse before age 16 | 1.66 | 1.09-2.53 | 0.017 |
| Forced first sexual intercourse | 1.42 | 1.00-2.02 | 0.047 |
| Woman's age: 25+ (ref:>25) | 1.54 | 1.09-2.17 | 0.015 |
| Not currently married (ref: married) | 1.96 | 1.35-2.83 | <0.001 |
| Woman has at least 8 years of education | 0.63 | 0.40-0.97 | 0.037 |
| Current partner is at least 30 years old | 1.43 | 1.02-2.01 | 0.037 |
| Past year intimate partner violence | 1.00 | 0.76-1.32 | 0.988 |

## Discussion

Our study highlights the prevalence of and association between childhood abuse experiences and subsequent HIV infection detected in antenxatal care. With the exception of a few small scale studies profiling corporal punishment in schools^6,26^we are not aware of any study in Zimbabwe on child sexual/physical abuse since 2000 that presents the prevalence of child sexual and physical abuse and association between childhood adversities and HIV positivity in antenatal care. Previous studies have found associations between more recent violent experiences or lifetime violent experiences against adult women^27^ without specifically studying the abuse that faced the women in childhood. Our study investigated a wide range of childhood abuse and/or conditions that have been associated with physical and sexual abuse such as a significantly older male partner experiences including sexual abuse, physical abuse, high age differences with a partner at first sex, forced first sexual intercourse and all were associated with HIV positivity. We failed to reject the hypothesis that childhood experiences of abuse are associated with HIV diagnosed in adulthood.

The prevalence of sexual abuse in children reported in this study falls within the prevalence range found in a systematic review of literature^10,28^ but is lower than found in African reviews^2^ where in some countries such as Egypt in which corporal punishment is legal as many as 50- 80% children reported physical punishment. The prevalence of physical abuse and sexual abuse is much lower than found in South Africa ^3,29^. The reported prevalence may have been a methodological underestimation of abuse experiences experienced by participants in a clinic population especially as we defined sexual abuse as any sexual intercourse occurring before the 15^th^ birthday when other studies define it until 16^th^ birthday. Research has shown that experiences of childhood violence among women is seldom exaggerated but can be under reported and underestimated. For example, some previous psychological experiments have shown that abusive experiences are forgotten if a person recently experienced similar and less abusive experiences in their lives than before^30^. Studies that interview children may give accurate prevalence figures although interviewing young people before they reach 18 would miss an opportunity to capture future violence before age 18.

We found a rate of FFS consistent with rates found in similar clinic based samples in Zimbabwe (11.8%)^21^, South Africa (12.4%) ^31^ and Uganda (14%) ^32^ but lower than reported in the Zimbabwe demographic and health survey (ZDHS) (21.6%)^33^ which collected data in the same year as in our study. While our study may have captured all childhood’s experiences, it may have suffered from recall bias. This might be because of the differences in study designs since the ZDHS collected population level data while our study was a clinic based study. Forced first sexual intercourse is a risk factor for a number of HIV risk factors such as IPV during pregnancy or after disclosing HIV status and severe revictimisation^31,34^ and as such prevention science must be concerned with such high prevalence to institute relevant behaviour change strategies.

We found that a significant proportion of women (14.3%) were sexually abused before age 15. Although this is lower than reported in South Africa it is of concern because early sexual debut increases risk of HIV infection among girls^21^. Young women who have sex early have a greater lifetime number of sexual partners and increased chances of partnering high risk men, usually older than them, have less education and less social integration^35^ than those who do not initiate sex early. Mathematical modelling with data in Zimbabwe has shown that a reduction in partnering with older partners reduces individual risk of HIV infection among women^36^. Therefore protecting children from abuse may be a viable entry point towards HIV prevention.

The analysis shows that both child sexual and physical abuse are major drivers of HIV risk as has been found in South Africa, also among low income women^29^.

The link between abusive events and HIV cannot be overemphasized. Proponents of the child development model argue that exposure to abuse in childhood negatively impacts on the child’s development leading to cognitive, psychological, and social impairment^12,37^. This may then lead to limited life skills consequently making them vulnerable to sexual abuse and HIV risk behaviours in relationships. Secondary victimisation in life is more likely in vulnerable people who have had the experience of violence and abuse normalised in their subjective experience ^31^. Some researchers have suggested two pathways linking violence in childhood with HIV infection. The first is a direct pathway that includes abusive experiences that lead to HIV infection through rape and forced sex. There is established evidence that points to the high likelihood of being HIV positive for people who rape girls and women^15,38–42^. Secondly, violence against women reduces one’s psychological wellbeing leading to risky behaviours and HIV infection as well as increasing fear and non-use of protective mechanisms that again lead to HIV infection^38^. Due to repeated exposure to violence one may lack the sense of urgency to resist unprotected sex which fuels HIV infection. Most of the discourse on child protection needs is contextualised more specifically in relation to orphans. However, our study suggests that child protection rights, including directed at non-orphans, must be discussed as part of HIV prevention strategies.

Despite significant findings, this study has several limitations that must be described to guide the interpretation of the results. Firstly, the study was cross sectional and involved a clinic population and as such does not allow making conclusions that childhood experiences lead to HIV infection. Even if we controlled for co-variates there may be other factors that we did not include in our questionnaire that may have contributed to such positive relationships. Prospective studies are required to identify possible causal relationships. Nevertheless, since we controlled for covariates, our results highlight the presence of the relationship between childhood abusive relationships and HIV. Secondly, asking women whose average age is 26 years about what happened before they turned 15 years may be subject to recall bias thereby underreporting the violence that happened to them^43^. Thirdly, some women may not report experiences that they perceive as minor or unimportant to report ^44,45^ or simply do not recall abusive events that happened many years back in their childhood^46^. However, studies have shown that abusive experiences are difficult to forget ^47^ and can haunt a person for a long time in their lives^48^ leading them to report them since a research setting presents an opportunity to vent their challenges and feel better. We did not measure psychosocial abuse in childhood, which is not easy, but emerging in studies on HIV and IPV. Lastly, our study only presented HIV status results and did not assess when the HIV was acquired as it is possible that they may have acquired it at the same time or prior to the abuse experiences. Our methodology has its strengths. Privately asking women about violence in primary health care settings offers a secure place to disclose painful experiences in one’s life compared to disclosing in household surveys where partners are usually available and limit women’s freedom to express themselves. A large sample size powered the study adequately to conduct multivariate analysis with a number of variables. Lastly, the analysis controlled for covariates and so helped to give a fuller picture of factors that are individually associated with HIV status.

## Conclusion

Exposure to child abuse is commonplace in Africa and happens in several ways highlighted in this study from physical to sexual violence, although this subject is not finding adequate attention from academic, implementing professionals and policy makers. The study therefore highlights the importance of protecting children as a way of preventing HIV infections in women and more broadly in society. Interventions to prevent HIV in women must therefore consider protecting children while researchers need to give priority to child abuse research in African settings.

## Declarations

### Author roles

SS conceived and designed the study and led the data collection, analysis and interpretation of data, drafted the article, led the revisions and approved the version to be published. NA, PS, MT, TS, CZ and substantially contributed towards study design, data analysis, and interpretation of data, revision of the manuscript and approved the final version to be published.

## References

1. Stoltenborgh, M., Bakermans-Kranenburg, M. J., Alink, L. R. A. & van IJzendoorn, M. H. The Prevalence of Child Maltreatment across the Globe: Review of a Series of Meta-Analyses. Child Abus. Rev. 24, 37–50 (2015).

2. Meinck, F., Cluver, L. D., Boyes, M. E. & Mhlongo, E. L. Risk and Protective Factors for Physical and Sexual Abuse of Children and Adolescents in Africa?: A Review and Implications for Practice. Trauma, Violence, Abus. 16, 81–107 (2015).

3. Shamu, S., Gevers, A., Mahlangu, B. P., Jama, P. N. & Chirwa, E. D. Prevalence and risk factors for intimate partner violence among Grade 8 learners in urban South Africa?: baseline analysis from the Skhokho Supporting Success cluster randomised controlled trial. (2015). doi:10.1093/inthealth/ihv068

4. Meinck, F. et al. Pathways From Family Disadvantage via Abusive Parenting and Caregiver Mental Health to Adolescent Health Risks in South Africa. J. Adolesc. Heal. 60, 57–64 (2017).

5. Meinck, F., Cluver, L.D., Boyes, M.E. Ndhlovu, L. D. Risk and protective factors for physical and emotional abuse victimisation amongst vulnerable children in South Africa. Child Abus. Rev. 24, 182–197 (2015).

6. Shumba, A. Epidemiology and etiology of reported cases of child physical abuse in Zimbabwean primary schools. Child Abuse Negl. 25, 265–277 (2001).

7. Carey, Paul D., Jennifer L. Walker, Wendy Rossouw, Soraya Seedat, D. J. S. Risk indicators and psychopathology in traumatised children and adolescents with a history of sexual abuse. Eur. Child Adolesc. Psychiatry 17, 93–98 (2008).

8. Jewkes, R. K., Dunkle, K., Nduna, M., Jama, P. N. & Puren, A. Associations between childhood adversity and depression, substance abuse and HIV and HSV2 incident infections in rural South African youth. Child Abuse Negl. 34, 833–41 (2010).

9. Birdthistle, I. J. et al. Child sexual abuse and links to HIV and orphanhood in urban Zimbabwe. J. Epidemiol. Community Health 65, 1075–1082 (2011).

10. Brown, D. W. et al. Exposure to physical and sexual violence and adverse health behaviours in African children: Results from the Global School-based Student Health Survey. Bulletin of the World Health Organization 87, 447–455 (2009).

11. Bensley, L. S., Van Eenwyk, J. & Simmons, K. W. Self-reported childhood sexual and physical abuse and adult HIV-risk behaviors and heavy drinking. Am. J. Prev. Med. 18, 151–158 (2000).

12. Norman, R. E. et al. The Long-Term Health Consequences of Child Physical Abuse, Emotional Abuse, and Neglect: A Systematic Review and Meta-Analysis. PLoS Medicine 9, (2012).

13. Lindert, J. et al. Sexual and physical abuse in childhood is associated with depression and anxiety over the life course: Systematic review and meta-analysis. International Journal of Public Health 59, 359–372 (2014).

14. Cohen, M. et al. Domestic violence and childhood sexual abuse in HIV-infected women and women at risk for HIV. Am. J. Public Health 90, 560–5 (2000).

15. Jewkes, R. et al. Factors associated with HIV sero-status in young rural South African women: Connections between intimate partner violence and HIV. Int. J. Epidemiol. 35, 1461–1468 (2006).

16. Gavin, L. et al. Factors Associated with HIV Infection in Adolescent Females in Zimbabwe. J. Adolesc. Heal. 39, (2006).

17. Schaefer, R. et al. Age-disparate relationships and HIV incidence in adolescent girls and young women: Evidence from Zimbabwe. AIDS 31, 1461–1470 (2017).

18. Shamu, S., Abrahams, N., Zarowsky, C., Shefer, T. & Temmerman, M. Intimate partner violence during pregnancy in Zimbabwe: A cross-sectional study of prevalence, predictors and associations with HIV. Trop. Med. Int. Heal. 18, 696–711 (2013).

19. Garcia-Moreno, C., Jansen, H. a F. M., Ellsberg, M., Heise, L. & Watts, C. H. WHO Multi-country Study on Women ’ s Health and Domestic Initial results on prevalence. Genetics 151, 277–83 (2005).

20. Dunkle, K. L. et al. Gender-based violence, relationship power, and risk of HIV infection in women attending antenatal clinics in South Africa. Lancet (London, England) 363, 1415–21 (2004).

21. Pettifor, A. E., Van Der Straten, A., Dunbar, M. S., Shiboski, S. C. & Padian, N. S. Early age of first sex: A risk factor for HIV infection among women in Zimbabwe. AIDS 18, 1435–1442 (2004).

22. Government of Zimbabwe. Sexual offences act. (2001).

23. Garcia-Moreno, C., Jansen, H. a F. M., Ellsberg, M., Heise, L. & Watts, C. H. Prevalence of intimate partner violence: findings from the WHO multi-country study on women’s health and domestic violence. Lancet 368, 1260–1269 (2006).

24. StataCorp. Stata Statistical Software: Release 13. 2013 (2013). doi:10.2307/2234838

25. World Health Organisation. Putting Women First: Ethical and safety recommendations for research on domestic violence against women. World Heal. Organ. 33 (2001).

26. Shumba, A., Mpofu, E., Chireshe, R. & Mapfumo, J. Corporal Punishment in Zimbabwean Schools?: Aetiology and Challenges. 1–10 (1995).

27. Widom, C. S., Czaja, S. & Dutton, M. A. Child abuse and neglect and intimate partner violence victimization and perpetration: A prospective investigation. Child Abuse Negl. 38, 650–663 (2014).

28. Barth, J., Bermetz, L., Heim, E., Trelle, S. & Tonia, T. The current prevalence of child sexual abuse worldwide: A systematic review and meta-analysis. Int. J. Public Health 58, 469–483 (2013).

29. Gibbs, A. et al. Childhood traumas as a risk factor for HIV-risk behaviours amongst young women and men living in urban informal settlements in South Africa: A cross-sectional study. PLoS One 13, e0195369 (2018).

30. Weems, C. F. et al. Memories of Traumatic Events in Childhood Fade After Experiencing Similar Less Stressful Events: Results From Two Natural Experiments. J. Exp. Psychol. Gen. 143, No Pagination Specified (2014).

31. Dunkle, K. L. et al. Prevalence and Patterns of Gender-based Violence and Revictimization among Women Attending Antenatal Clinics in Soweto, South Africa. 160, 230–239 (2004).

32. Koenig, M. a et al. Coerced first intercourse and reproductive health among adolescent women in Rakai, Uganda. Int. Fam. Plan. Perspect. 30, 156–163 (2004).

33. Zimbabwe National Statistics Agency, I. C. F. I. I. Zimbabwe Demographic and Health Survey 2010-11. 1–470 (2012).

34. Shamu, S. The dynamics of intimate partner violence during pregnancy and linkages with hiv infection and disclosure in zimbabwe. (International Centre for Reproductive Health, 2013).

35. Hallett, T. B. et al. Age at first sex and HIV infection in rural Zimbabwe. Stud. Fam. Plann. 38, 1–10 (2007).

36. Hallett, T. B., Gregson, S., Lewis, J. J. C., Lopman, B. A. & Garnett, G. P. Behaviour change in generalised HIV epidemics: Impact of reducing cross-generational sex and delaying age at sexual debut. Sex. Transm. Infect. 83, (2007).

37. Perry, B. D. The Neurodevelopmental Impact of Violence in Childhood. Textb. Child Adolesc. Forensic Psychiatry 221–238 (2001).

38. Dunkle, K. L. & Decker, M. R. Gender-Based Violence and HIV: Reviewing the Evidence for Links and Causal Pathways in the General Population and High-risk Groups. American Journal of Reproductive Immunology 69, 20–26 (2013).

39. Jewkes, R. et al. A cluster randomized-controlled trial to determine the effectiveness of Stepping Stones in preventing HIV infections and promoting safer sexual behaviour amongst youth in the rural Eastern Cape, South Africa: Trial design, methods and baseline findings. Trop. Med. Int. Heal. 11, 3–16 (2006).

40. Khanna, A., Goyal, R. S. & Bhawsar, R. Menstrual Practices and Reproductive Problems: A Study of Adolescent Girls in Rajasthan. J. Health Manag. 7, 91–107 (2005).

41. Raj, A. et al. Perpetration of Intimate Partner Violence Associated With Sexual Risk Behaviors Among Young Adult Men. 96, 1873–1878 (2006).

42. Silverman, J. G., Decker, M. R., Kapur, N. A., Gupta, J. & Raj, A. Violence against wives, sexual risk and sexually transmitted infection among Bangladeshi men. 211–215 (2007). doi:10.1136/sti.2006.023366

43. Williams, L. W. Recall of childhood trauma: A prospective study of women’s memories of child sexual abuse. J. Consult. Clin. Psychol. 62, 1167–1176 (1994).

44. Pallitto, C. C. et al. Intimate partner violence, abortion, and unintended pregnancy: results from the WHO Multi-country Study on Women’s Health and Domestic Violence. Int. J. Gynaecol. Obstet. 120, 3–9 (2013).

45. Gil-González, D., Vives-Cases, C., Ruiz, M. T., Carrasco-Portiño, M. & Álvarez-Dardet, C. Childhood experiences of violence in perpetrators as a risk factor of intimate partner violence: A systematic review. J. Public Health (Bangkok). 30, 14–22 (2008).

46. Williams, L. M. Recall of childhood trauma: A prospective study of women’s memories of child sexual abuse. J. Consult. Clin. Psychol. 62, 1167–1176 (1994).

47. Bernet, C.Z.; Stein, M. B. Relationship of childhood maltreatment to the onset and course of major depression in adulthood. Depress. Anxiety 9, 169–174 (1999).

48. Schwab, G. Haunting legacies: Violent histories and transgenerational trauma.,. (Columbia University Press, 2010).

